# Grazing impacts on ground beetle (Coleoptera: Carabidae) abundance and diversity on a semi-natural grassland

**DOI:** 10.1101/2020.08.29.273797

**Authors:** Gabor Pozsgai, Luis Quinzo-Ortega, Nick A. Littlewood

**Affiliations:** The James Hutton Institute, Craigiebuckler, Aberdeen, AB15 8QH, UK; Department of Applied Ecology, Fujian Agriculture and Forestry University, 15 Shangxiadian Road, Cangshan District, Fuzhou, Fujian, 350002, China; School of Biological and Environmental Sciences, James Parsons Building, Liverpool John Moores University, Byrom Street, Liverpool L3 3AF; Department of Rural Land Use, SRUC North Facility, SRUC Aberdeen, Ferguson Building, Craibstone Estate, Bucksburn, Aberdeen AB2l 9YA, UK

**Keywords:** Grazing pressure, insect assemblages, livestock, sustainable habitat management, upland grassland

## Abstract

Semi-natural grasslands are commonly managed as a grazing resource for domestic livestock but, due to their unique biodiversity, they are also of conservation interest. Numerous drivers have impacted on the status of these grasslands in recent decades, most importantly changing grazing management strategies. These changes have the potential to affect the biodiversity associated with these habitats, including on some rich invertebrate assemblages. Responses, however, are often dissimilar between different invertebrate taxa.

We investigated the responses of ground beetles to different grazing regimes within a long-term grazing experiment on upland semi-natural grassland in Scotland. Although there was substantial overlap between ground beetle assemblages in different grazing treatments, species richness, mean abundance and Shannon diversity of ground beetles were significantly lower in ungrazed plots than in plots subject to high- or low-intensity sheep grazing. Ground beetle abundance (but not species richness or diversity) were lower in ungrazed plots compared to those with low-intensity mixed grazing by sheep and cattle. However, no differences were identified in abundance, species richness or diversity between the three grazed treatments.

Our results suggest that ground beetles may show different responses to grazing compared to responses of some other invertebrate groups and demonstrates the difficulty of carrying out management for a multi-taxon benefit.

## 1 Introduction

Semi-natural grasslands around the world are valued for their unique biodiversity (e.g. WallisDeVries et al., 2002). Recently, they have also been recognised as being important carbon stores (e.g. Conant et al., 2017). The nature of these grasslands, which have not suffered deleterious effects of fertilizer or herbicide application, varies with geographic location and factors such as altitude. However, semi-natural grasslands from across their range are under threat (e.g. Oakleaf et al., 2015; Ridding et al., 2015), with drivers of change including altered livestock grazing, land use changes (Török et al., 2016), biological invasions, and wildfires (Coates et al., 2016).

Among semi-natural grasslands that have a long-term history of grazing by domestic herbivores, the condition of upland grasslands areas has gained considerable attention in recent decades (Evans et al., 2015; McGovern et al., 2011). In Europe, for example, recent changes in agricultural policy have driven substantial shifts in the way that semi-natural upland habitats are managed. In some areas, such as in the UK, this has manifested in the number of grazing livestock declining markedly since the start of the twentieth century and many areas that were formerly grazed now have no grazing by domestic herbivores (e.g. Martin et al., 2013). Resultant changes in vegetation structure and composition may be slow to become apparent in these low-productivity systems (Pakeman et al., 2019) and may vary with livestock type (Tóth et al., 2018) but evidence is gradually emerging that such land-use changes are having a marked impact on a range of biodiversity, including birds (Evans et al., 2015) and invertebrates(Dennis et al., 2007). The extent of such impacts, however, varies widely amongst different invertebrate groups (Bonari et al., 2017; García et al., 2009).

Ground beetles (Coleoptera: Carabidae) populations have suffered large scale declines in the UK over recent decades(Brooks et al., 2012; Pozsgai and Littlewood, 2014) and increased homogeneity of assemblages (Pozsgai et al., 2015) at systematically monitored sites. The causes of these declines are not well understood. Some declines may be linked to agricultural intensification, including, for example, improvement of grassland for grazing animals (Luff and Rushton, 1989). Evidence of detrimental effects of climate change is also growing. For example, changing phenologies of some ground beetle species have been linked with climate change (Pozsgai et al., 2018; Pozsgai and Littlewood, 2014, 2011) and changes in upland ground beetle assemblages and an increased prevalence of generalist species have been linked to decreasing maximum temperatures and increasing rainfall (Pozsgai et al., 2015).

The extent to which differences in grazing management may impact on ground beetle abundances and assemblages has been studied in upland heathland(Gardner et al., 1997) and in upland calcareous grassland (Lyons et al., 2017) but rather less so in acidic grasslands that dominate much of the upland vegetation in the UK (though see Cole et al., 2006). There is some evidence from elsewhere that ground beetle abundance and, especially, species richness may be reduced by relaxation or cessation of grazing (e.g. Grandchamp et al., 2005). However, such trends may be site and context specific with, for example, Twardowski et al. (2017) reporting more ground beetles in sites that were less intensively grazed.

Ground beetles are among the more common of surface feeding arthropods and have important functional roles to play in grasslands as predators of smaller invertebrates (e.g. Lövei and Sunderland, 1996) in addition to being prey items themselves, including for upland birds (Buchanan et al., 2006). Given the reported declines in ground beetle abundances and ongoing changes in upland management, it is desirable to understand what forms of management may best support populations. To investigate the influence of the intensity of one aspect of management, livestock grazing, on ground beetle assemblages of semi-natural upland acidic grassland, we sampled ground beetles within a replicated, controlled grazing experiment. In particular, we hypothesised that, as reduced grazing leads to increased vegetation density, which may impede ground beetle foraging ability, the abundance (as measured by activity density), species richness and diversity of ground beetles would be positively correlated with grazing intensity. Furthermore, we assessed whether or not any such differences that we identified between grazing treatments resulted in differences in the overall ground beetle assemblage structure.

## 2 Methods

### 2.1 Field Site

Sampling was carried out on the estate of Glen Finglas, in Perthshire, Scotland (56°16’N, 4°24’W). This estate is owned and managed by a conservation charity, the Woodland Trust, with a long-term aim of restoring a semi-natural woodland/grassland mosaic. It extends to 4,085 ha in area. Large parts of the estate remain open ground, dominated by acid grassland and mire. The most represented National Vegetation Classification (NVC) communities (Rodwell, 1992, 1991) in the study areas were M23 (*Juncus effusus/acutiflorus–Galium palustre* rush-pasture), M25 (*Molinia caerulea–Potentilla erecta* mire), U4 (*Festuca ovina–Agrostis capillaris–Galium saxatile* grassland) and U5 (*Nardus stricta–Galium saxatile* grassland). Some areas were covered by bracken (*Pteridium aquilinum*, NVC U20). Sample sites ranged in altitude from approximately 200 m to 500 m.

### 2.2 Treatments

Six replicate blocks (labelled A to F) of four grazing treatment plots were established in 2003. Each plot measured 3.3 ha. Plots were located at three sites, each containing two replicate blocks of each treatment (so a total of eight plots in each of three sites). There sites were each separated by approximately 5 km. The grazing treatments were high-intensity sheep grazing, with nine sheep per plot (Treatment I), low-intensity sheep grazing with three sheep per plot (Treatment II), low-intensity mixed grazing with two sheep per plot and, for four weeks in August/September, two cows per plot, each with a suckling calf (Treatment III) and ungrazed by domestic herbivores (Treatment IV). Sheep were removed from plots during severe weather over winter and for routine farming practices, such as shearing. Grazing treatments were assigned randomly at the outset across the four plots for each block (Figure 1).

**Figure 1:**
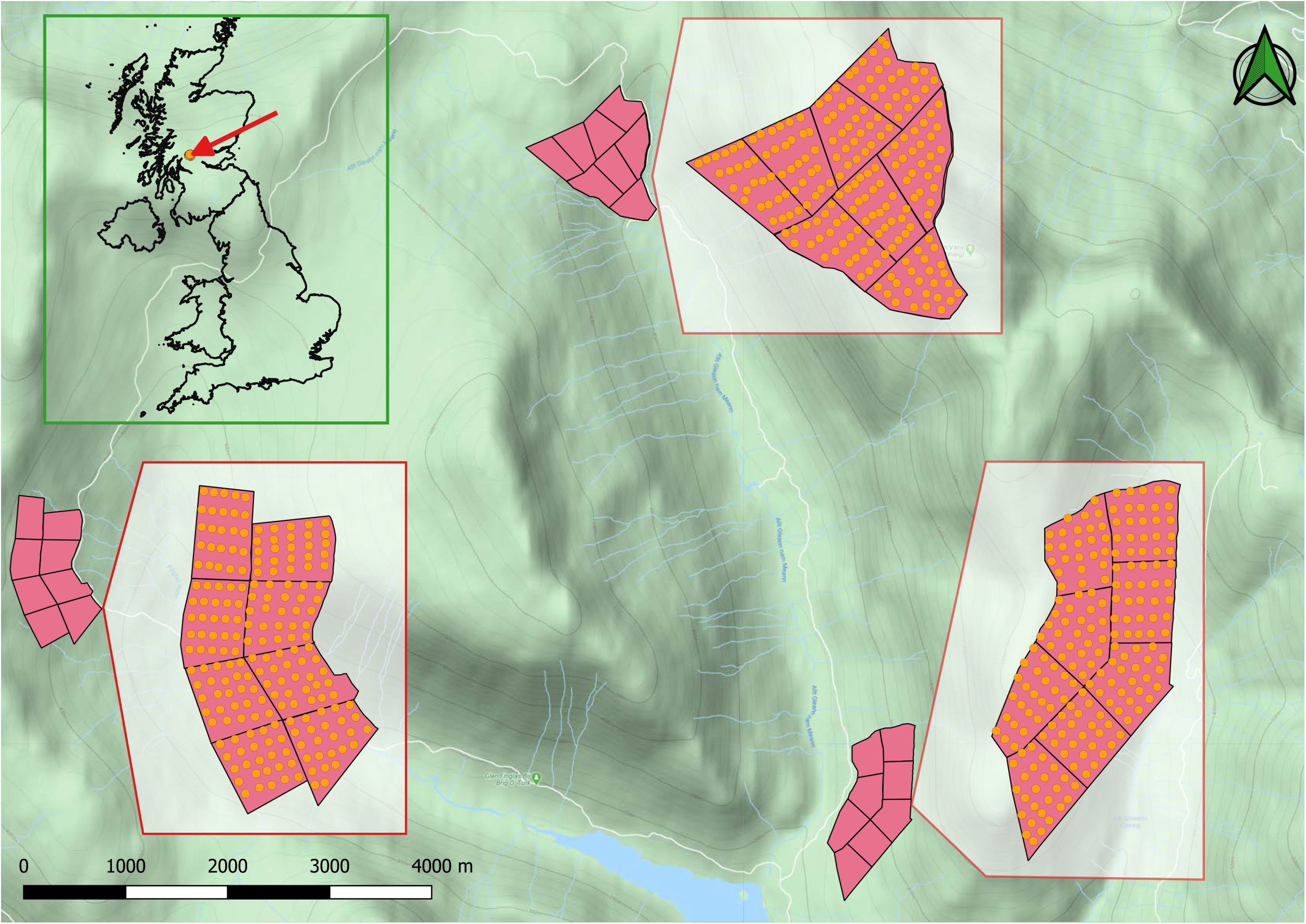
Location of sampling points at Glen Finglas, Scotland. The overview map in the topleft corner shows the location within the UK, the main map shows the arrangement of sample blocks within the glen and the larger scale maps show the arrangement of sample points within these blocks.

### 2.3 Ground beetle sampling

Sampling was carried out from 2009 to 2013. Each year, five points were chosen randomly from a pre-established grid of 25 sampling points within each plot. A 500-ml cup, with a diameter of 8 cm, was used as a pitfall trap and was placed within 1 m of each of these five points. Each cup was part-filled with ethylene-glycol that served as a preservative. Chicken wire, with a mesh size of 2 cm, was attached over the top of each trap to minimise capture of small vertebrates and to deter interference by sheep and a chicken wire dome was placed in each cup to aid escape of any vertebrates that did fall in the trap. Traps were set in May each year and emptied and reset at approximately three-week intervals through to September. Trap contents were stored in a freezer until sorting and identification in the laboratory. Subsequently, they were transferred to 70% ethanol and stored at the James Hutton Institute, Aberdeen, UK. Specimens were identified to species by reference primarily to(Lindroth, 1986, 1985) and Luff (2007). Species taxonomy follows Luff (2007).

### 2.4 Data analysis

The mean number of each ground beetle species captured was calculated across sampling dates within a year, resulting in a data matrix showing sampling point/year/treatment/block in rows and species in columns. Commonly used diversity measures, namely abundance and species richness per trap, and Shannon-Wiener diversity were calculated, and differences between treatments were investigated using Kruskal-Wallis test. When significant differences were found, post-hoc pairwise Wilcoxon tests were conducted to compare individual treatments. In order to address multiple comparison issues, p-values were corrected using Holm’s method.

Non-metric multidimensional scaling (NMDS) was used on a Bray-Curtis distance matrix to show the similarities of species composition among treatments and to visualize multivariate patterns in two dimensions. Permutational Multivariate Analysis of Variance Using Distance Matrices (ADONIS), implemented in the vegan R package (Oksanen et al., 2010), was used to detect the main drivers of ground beetle assemblage composition, with treatment as a fixed variable and sampling block and sampling year as grouping factors. *P*-values were calculated through a permutation process with 999 iterations. Species that were the most important in discriminating between treatments were selected using similarity percentages (Clarke, 1993) with the help of the simper() function in the vegan package. Indicator value analysis, IndVal (De Cáceres et al., 2010; De Cáceres and Legendre, 2009), was used to investigate whether any specific species were strongly associated with particular treatments, experimental blocks or years. Prior to the multivariate analysis, ground beetle numbers were transformed using Hellinger’s method as suggested by O’Hara and Kotze (2010).

The impact of grazing pressure on the activity-density of the seven most common ground beetle species was analysed using linear mixed effect models, where grazing treatment and sampling year were considered as fixed, and sampling block as random effects. Sampling year was considered as a fixed effect because climate change (Pozsgai et al., 2015) and altered land use (Déri et al., 2011; York, 2000)can cause a directional shift in insect assemblages over time. Thus, temporal trends in the abundance of species in our samples were also expected. Moreover, these abundance trends could also be of ecological interest (e.g. common species declining with less common, habitat specialists, increasing) which further supports the inclusion of sampling year amongst the fixed effects.

All analyses were carried out with the R statistical software, using vegan (Oksanen et al., 2010), indicspecies (De Cáceres and Legendre, 2009), and nlme (Pinheiro et al., 2017) packages.

## 3 Results

Thirty-six species of ground beetle were recorded (Figure 2). These comprised 5,120 individual ground beetles (Supplementary Material 1). The most frequently caught species were *Pterostichus nigrita* (1,217 individuals), *Pterostichus madidus* (1,050 individuals) and *Pterostichus niger* (545 individuals). Thirteen species were represented by fewer than ten individuals each. The mean number of beetles caught per trap per collection was relatively low, at 7.02 ± 7.00 (mean ± IQR).

**Figure 2:**
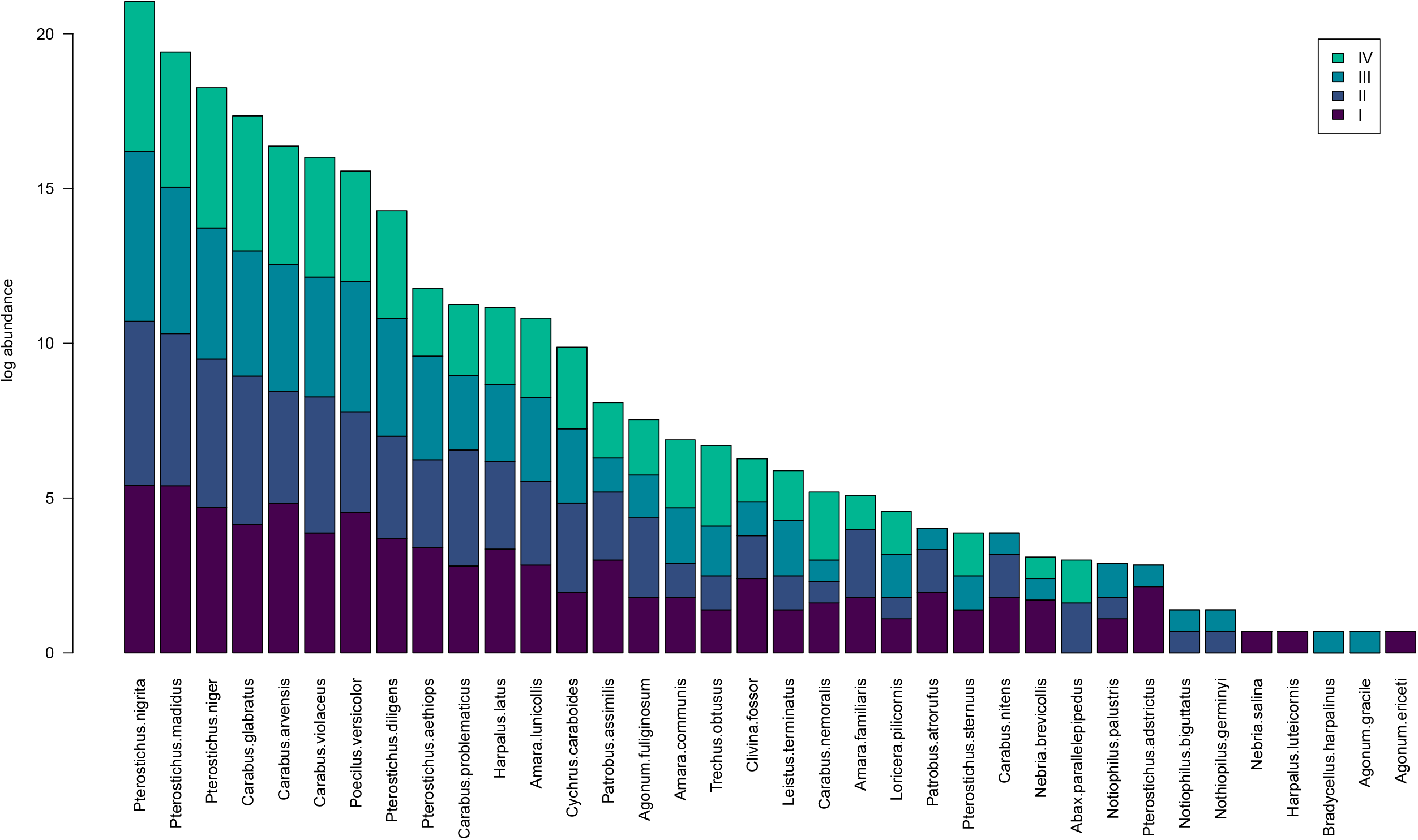
Ground beetle species recorded at Glen Finglas, 2009-2013, showing log abundance in each grazing treatment.

### 3.1 Ground beetle abundance and diversity

Species richness, mean abundance and Shannon diversity were significantly lower in treatment IV (ungrazed) than in treatments I (high-intensity sheep) and II (low-intensity sheep). Both species richness and Shannon diversity were greater in treatment I than in III (low intensity sheep and cows) and mean abundance was significantly higher in treatment III than in treatment IV (Figure 3). There were no significant overall temporal trends in mean abundances and species richness but some individual plots showed signs of decline in species richness (Supplementary Material 2).

**Figure 3:**
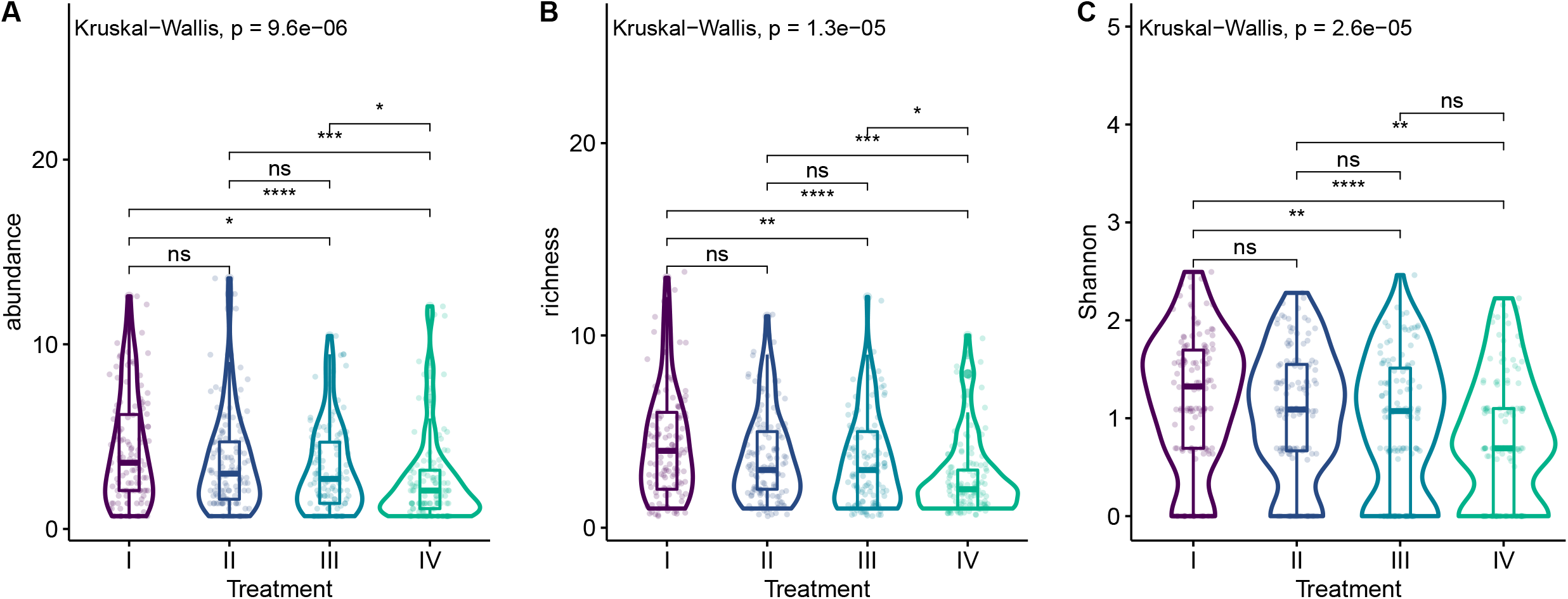
The effect of grazing treatments on ground beetle diversities. Violin plots show probability densities of (A) log-transformed abundance, (B) species richness, and (C) Shannon diversity. Boxplots inside violins represent the distribution of measured data and show the median, and the lower and upper quartiles. Individual data points are also displayed. Results of pairwise comparisons are indicated on the top of the lines connecting violins. Adjusted p-values are indicated as *: *p* < 0.05; **: *p* < 0.01; ***: *p* < 0.001; NS: not significant (*p* > 0.05).

### 3.2 Ground beetle assemblages

Multidimensional scaling showed that there was a large overlap between ground beetle assemblages in the four grazing treatments (Figure 3). The ADONIS model, however, demonstrated a significant treatment effect, along with significant effects from sampling year, sampling block, and the combination of these (Table 1). The ADONIS model explained 36.3% of the total variance, of which the most important was the block effect (7.4%). Only a small proportion of variance, 2.3%, was explained purely by the treatment effect. Pairwise ADONIS showed a significant difference in assemblages between treatment I and the ungrazed treatment IV (F = 3.98, R^2^ = 0.064, adjusted *p*-value = 0.006) but none of the other treatments were significantly different in pairwise comparison. When data were separated to blocks, the treatment effect was only significant in three of the six blocks, one from each block pair. The difference was mostly caused by treatment IV, which was well separated from all other treatments. Treatments I, II and III showed little separation from each other (Figure 4).

**Table 1:** The effects of treatment, sampling block, and sampling year on carabid assemblages. The results of the Permutational Multivariate Analysis of Variance Using Distance Matrices (ADONIS) on species assemblage distance matrix. df: degrees of freedom, SS: sum of squares.

|  | Df | SS | F-value | p-value |
| --- | --- | --- | --- | --- |
| **Treatment** | 3 | 1.002 | 2.946 | 0.001 |
| **Year** | 4 | 1.029 | 2.271 | 0.001 |
| **Block** | 5 | 4.062 | 7.168 | 0.001 |
| **Treatment x Year** | 12 | 1.428 | 1.050 | 0.377 |
| **Treatment x Year** | 15 | 2.611 | 1.536 | 0.004 |
| **Residual** | 78 | 8.840 |  |  |

**Figure 4:**
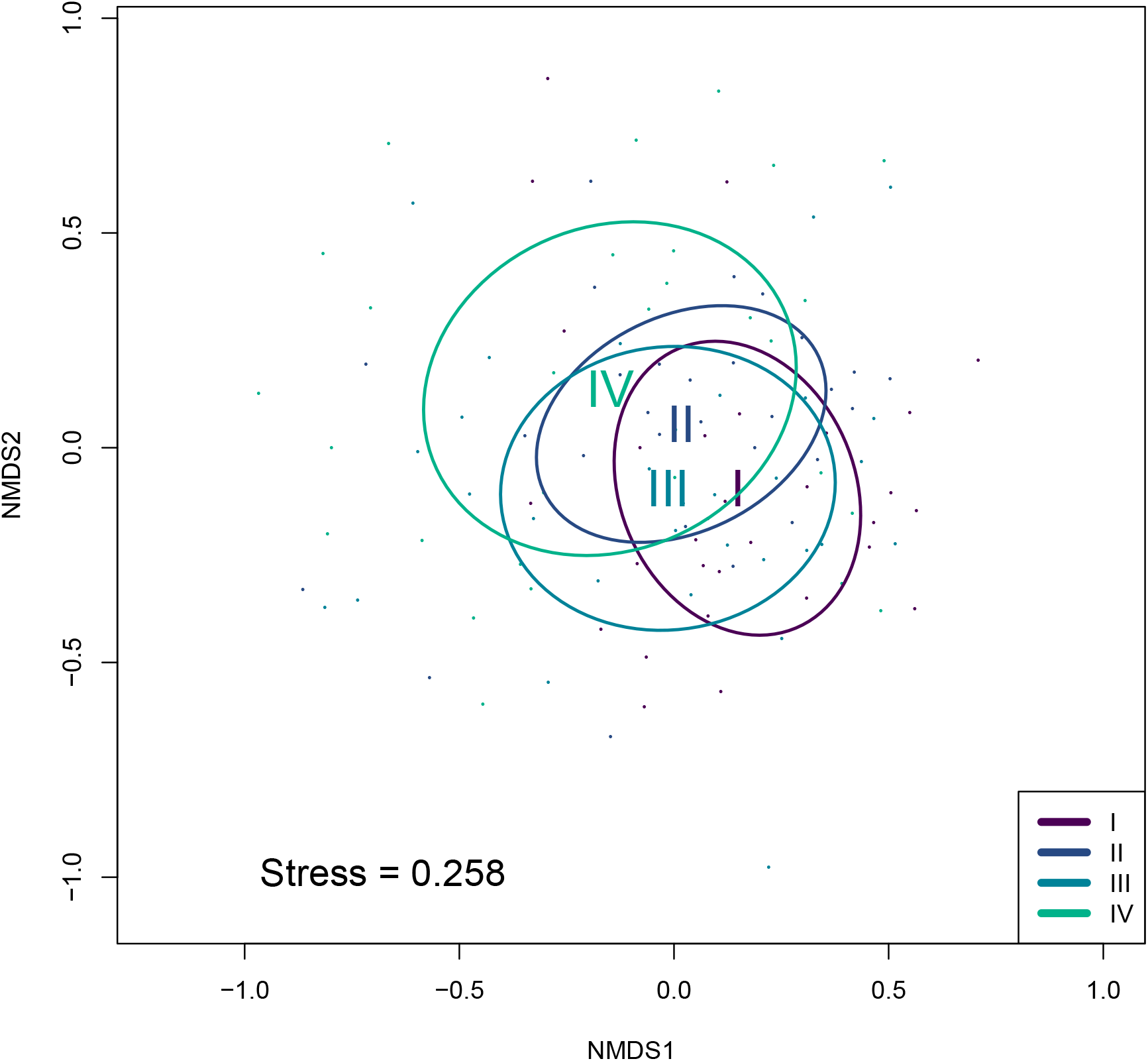
Multidimensional scaling of ground beetle samples from Glen Finglas Numerals I, II, III and IV show grazing treatments and are positioned precisely centrally relative to the ellipse for that treatment. Ellipses indicate areas within which 95% of the site points belonging to one treatment are located.

**Figure 5:**
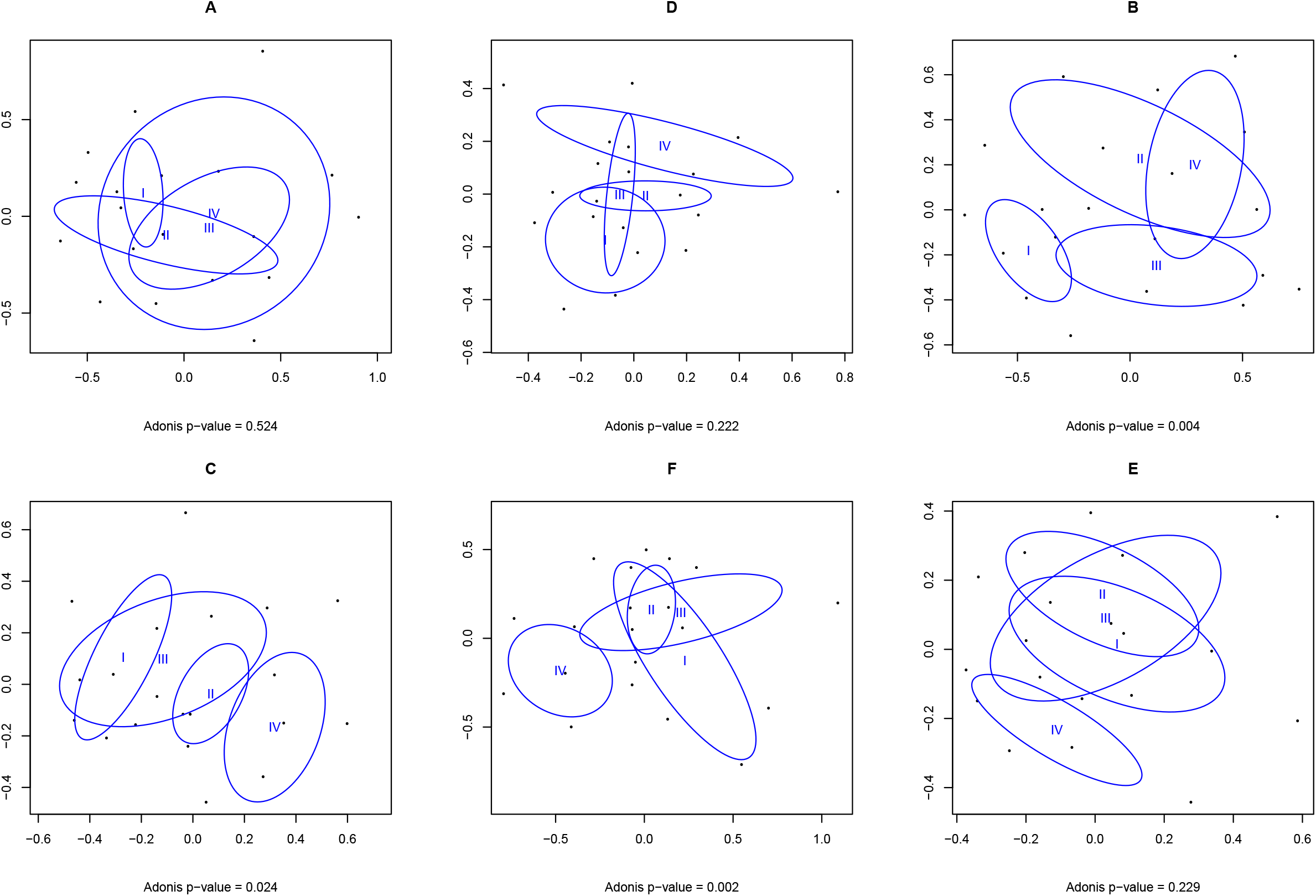
Multidimensional scaling of ground beetle samples from Glen Finglas. Each plot, A to E, represents a treatment block. Numerals I, II, III and IV show grazing treatments and are positioned precisely centrally relative to the ellipse for that treatment. Ellipses indicate areas within which 95% of the site points belonging to one treatment are located.

Although ADONIS indicated a significant effect of sampling year, no temporal trends were apparent in the assemblages overall (Supplementary Material 3). Data suggested, though, that assemblages in treatments II, III and IV were changing over time in some treatment blocks. In particular the ecological distance between samples increased over the years within treatment block B, and also treatment IV of block A and in treatment III of block F. The ecological distance between treatments I and IV showed an increase only in block E. Simper analysis indicated that the two most important species in shaping differences in ground beetle assemblages between treatments were *Pterostichus nigrita* and *Pterostichus madidus*. Further species played important roles in shaping differences between assemblages, notably *Pterostichus niger* (in differences between treatments I and II and between I and IV), *Carabus arvensis* (differences between treatments I and III) and *Carabus glabratus* (differences between treatments II and III, treatments II and IV and treatments III and IV) (Supplementary material 4).

Common, habitat generalist species dominated all treatments. Among the species that are commonly associated with wet peatlands (e.g. *Agonum ericeti, Patrobus atrorufus, Trechus obtusus* (Blake et al., 2003) only a few that are also tolerant of drier soils, such as *T. obtusus* (Eyre et al., 1989) were found in the ungrazed treatment IV plots and, indeed, IndVal analysis pinpointed this species as an indicator for treatment IV (IndVal g = 0.231, *p* = 0.012). In contrast, *Pterostichus adstrictus* (IndVal g = 0.224, *p* = 0.001) was typically found in the most heavily grazed treatment I. No other species were identified by this analysis as being significant indicators of any single grazing treatment (Table 2).

**Table 2:**
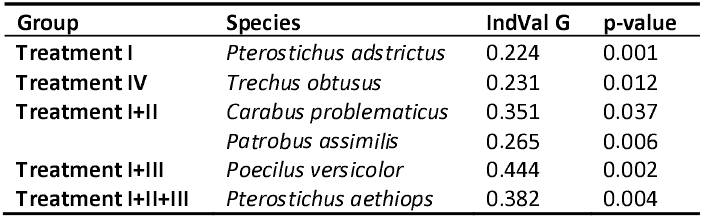
Indicator values and bootstrapped p-values of ground beetle species with high fidelity to treatments or treatment groups.

| Group | Species | IndVal G | p-value |
| --- | --- | --- | --- |
| **Treatment I** | *Pterostichus adstrictus* | 0.224 | 0.001 |
| **Treatment IV** | *Trechus obtusus* | 0.231 | 0.012 |
| **Treatment I+II** | *Carabus problematicus* | 0.351 | 0.037 |
|  | *Patrobus assimilis* | 0.265 | 0.006 |
| **Treatment I+III** | *Poecilus versicolor* | 0.444 | 0.002 |
| **Treatment I+II+III** | *Pterostichus aethiops* | 0.382 | 0.004 |

The regression of the mean abundances of the eight most commonly caught species showed a significant treatment effect (Block was included as a random effect) only for *Carabus arvensis* (*p* = 0.042). Sampling year caused a significant difference in abundance between treatments for *Pterostichus nigrita* (*p* = 0.002), Carabus arvensis (*p* = 0.048) and *Pterostichus diligens* (*p* = 0.001). Differences, however, were not significant for *C. arvensis*, when years were pairwise compared and the Tukey HSK correction was applied (Supplementary material 5).

## 4 Discussion

Overall, there was substantial overlap between ground beetle assemblages in different grazing treatments. We did find that there were significant differences between treatments, though these were small, with treatment explaining just a small portion of the variance in the complete dataset. Nonetheless, in partial support of our hypothesis, our results also showed that there were fewer ground beetles caught, fewer species recorded and lower diversity in the ungrazed treatment IV than in the two sheep-grazed treatments (treatments I and II) and that fewer were caught in treatment IV than in the mixed grazing treatment III. However, no differences were identified in abundance, species richness or diversity between the three grazed treatments.

### 4.1 Grazing impact on ground beetle abundance

The finding that ground beetles were caught in lower overall numbers in ungrazed grassland is broadly consistent with results from several other studies from a variety of grassland habitats. For example, ground beetle abundance was higher in grazed compared to ungrazed sites in three different dune grassland types in the Netherland (Nijssen et al., 2001). Similarly, Pétillon et al. (2007) reported higher ground beetle abundance (and species richness) in areas of saltmarsh in France that were subject to sheep grazing and grass cutting than in non-grazed and uncut areas, and this trend was driven by typical grassland species. In a study in Lapland ground beetles were more numerous on plots grazed by reindeer *Rangifer tarandus* than on ungrazed plots but their numbers peaked under intermediate grazing levels (Suominen et al., 2003), suggesting that there may be an optimum level of grazing at which ground beetle abundance is maximised.

This effect of grazing may be replicated by other forms of vegetation removal. For example, Gimingham (1985) showed that ground beetles were most abundant in the pioneer (youngest) stage of growth of the dwarf shrub, *Calluna vulgaris,* (under a rotational burning regime), when vegetation complexity was at its lowest. In parallel, Sanderson et al. (2020) found more ground beetles in areas of *C. vulgaris* that were mechanically cut between one and seven years previously compared to areas cut eight or more years previously. Although dwarf shrubs were not dominant in our study plots, these results are consistent with our finding that ground beetle activity density was lower in the ungrazed treatment, which had the tallest vegetation with the greatest biomass (Evans et al., 2015) and most extensive areas of dense grassy tussocks (Smith et al., 2014), than in the grazed treatments. This observation may be at least partly driven by the two species that occurred in the highest numbers in our study, namely *Pterostichus nigrita* and *Pterostichus madidus*. These have previously been shown to display trends of decreasing abundance with increasing vegetation height (e.g. Dennis et al., 1997). Thus, the response of these common species may drive the higher overall abundances in plots that were grazed, and thus had shorter vegetation(Pakeman et al., 2019), than in the ungrazed plots.

Assessments of ground beetle abundance using pitfall traps can lead to bias in that more mobile species are more likely to encounter, and be caught by, the traps (e.g. Brown and Matthews, 2016) whilst captures may also be biased by ground beetle body size (Hancock and Legg, 2011), perhaps due to correlation between size and mobility. Thus, the results represent “activity density” rather than indices of absolute abundance. If mobility is reduced in denser vegetation, this might cause an apparent reduction in the number of individuals caught in our ungrazed treatment and an apparent increase in those in the highest grazing intensity treatment. Such an effect has been previously observed in an experimental set-up when there was no evidence of differences in absolute ground beetle density between areas of different vegetation structure(Thomas et al., 2006). Other studies based on pitfall trap data have similarly found lower activity density in areas of denser vegetation (e.g. Cole et al., 2008). However, as activity density cannot be directly translated into abundance, we are unable to demonstrate whether such a difference represents an actual abundance difference. Nonetheless our samples were taken using a sufficiently large trap and over sufficient timescales to meet requirements recommended by Jung et al. (2019) for promoting reliability of pitfall trapping programmes.

### 4.2 Grazing impact on ground beetle diversity

Similarly to abundance, species richness of ground beetles has been shown in some studies to be enhanced by grazing. For example, in montane meadows in Switzerland, increased density of cattle grazing was associated with increased ground beetle species richness (Grandchamp et al., 2005) and, in Sweden, species richness was positively (albeit weakly) associated with increased cattle grazing intensity (Söderström et al., 2001). However, some studies have found no significant relationship, including on semi-natural grasslands in Hungary, where ground beetle species richness was not affected by grazing intensity of cattle (Batáry et al., 2007) and a similar lack of significant impact was reported from British Columbia, Canada (Bassett and Fraser, 2015). An opposite finding to ours was that, on heather moorland in Scotland, ground beetle species richness was reduced on sites that were the most heavily grazed by deer and sheep (Gardner et al., 1997). It is possible in this latter case that the highest intensity sheep-grazing treatment was higher, in terms of the quantity of forage that remained after grazing, than in our high-intensity sheep-grazing treatment and that more discrimination between treatments may have been apparent had we used a higher sheep stocking rate.

Metrics of ground beetle diversity may also be positively correlated to vegetation removal. Grandchamp et al. (2005), for example, found that intensive management of grasslands (through mowing, grazing and fertilisation) helped to maintain ground beetle assemblages with higher diversity than of those in grasslands managed at a lower intensity. Similarly, rotational burning, which reduces biomass of *C. vulgaris,* may enhance ground beetle diversity (Gardner et al., 1991).

### 4.3 Grazing impact on ground beetle assemblage structure

Although, as discussed above, variation in species mobility may affect ground beetle captures, we utilised a substantial number of traps set across an extensive experimental site. This was on a scale reported to be sufficient to accurately sample the assemblage that was present (Lövei and Magura, 2011). This is further evidenced by the ground beetle assemblages at our site being broadly similar in the different treatments. Additionally, the fact that Simper analysis indicated that the most abundant species were responsible for the significant differences that we did identify between assemblages suggests that those differences were more quantitative rather than qualitative. Previously, work has documented distinct ground beetle assemblages associated with upland environments in the UK (e.g. Luff et al., 1992) and shown that these differ between different upland habitats (Butterfield and Coulson, 1983). In particular, ground beetle assemblages have been shown to be substantially shaped by environmental conditions, including both vegetation structure and soil conditions (e.g. Luff et al., 1992, 1989). In comparable habitat to that in our study, Cole et al. (2006) found that large flightless *Carabus* species favoured an extensive grazing regime over an intensive one. However, other research has failed to find species that are indicators or different management treatments within habitat mosaics (e.g. Pravia et al., 2019). This is in line with our own findings in which we did not identify species or assemblages that were particularly associated with different intensities of grazing at our site, including no evidence that some species preferred the less intensively grazed plots that might equate to a more extensive grazing regime.

Inconsistent responses to grazing apparent among ground beetle assemblages(Bonari et al., 2017; García et al., 2009; Lengyel et al., 2016), suggest that this group may not be suitable as an indicator taxon for grassland management, including in restoration projects. Lengyel et al. (2016), in their multi-taxon study, pointed out clear differences in the ways that a range of organism groups respond to vegetation diversity and structure. Some may more consistently show directional responses to treatment. For example, plant communities, appear to be positively affected by grazing (Török et al., 2014). For ground beetles, although Koivula (2011) showed that European ground beetles may reflect management, the requirement for further research into the usefulness of these responses for conservation purposes was stressed and our own results suggest that their use as indicators may be limited, at least within the system that we studied.

### 4.4 Implications for upland grassland management

Previous studies, based on sampling within the same long-term experiment as this work, have shown a range of responses of invertebrate faunal groups to the four grazing treatments. Foliar invertebrates as a whole have been shown to respond positively to relaxation of grazing levels and cessation of grazing, over the first three years following the application of grazing treatments (Dennis et al., 2007). Studies of both moths (Littlewood, 2008) and Auchenorrhyncha (Littlewood et al., 2012), sampled five years on from the initiation of grazing treatments, likewise showed trends to greater abundance and species richness in less grazed compared to more grazed plots. In both cases, abundance and species richness were significantly higher in the ungrazed treatment plots compared with the high-intensity sheep grazing treatment. Overall arthropod abundance, measured at intervals over the first nine years of the experiment, was also negatively related to grazing intensity (Evans et al., 2015). Our results show only small differences between treatments but those that were identified contrasted with these previous results by showing the lowest abundance and species richness in ungrazed plots. This might suggest that whilst potential prey items for carabids may be more abundant with reduction in grazing pressures, ground beetles are not able to fully exploit this increased resource availability, perhaps due to those prey being less accessible in the denser vegetation. This demonstrate how environmental factors act quite differently compared to other taxa in shaping occurrences within this largely carnivorous group and indicates that management for a multi-taxon benefit may be difficult to achieve on sites under a single uniform management regime (Kruess and Tscharntke, 2002).

Multi-species livestock farming has been shown to benefit biodiversity (Martin et al., 2020), and also to promote ecosystem multifunctionality (Wang et al., 2019). Our results on carabid assemblages, however, were not supportive to these studies; abundances, species richnesses, as well as Shannon diversities were lower in plots grazed by a mix of cattle and sheep than in those with high-intensity sheep grazing and showed no difference from other treatments. Whereas these results highlight that optimising upland grazing for multi-taxon benefits with mixed-species grazing is unlikely to equally favour all species, knowing which taxa are ‘winners’ or ‘losers’ is essential for successfully managing diverse grasslands.

## 5 Conclusions

In this study, we demonstrate that ground beetle assemblages are affected by differences in livestock grazing in our upland experimental site. In particular, abundance, species richness and diversity of ground beetles were lower in the ungrazed treatment than in the high- or low-intensity sheep grazed treatments. These results contrast with previous studies of other invertebrate taxa and show, in particular, that a single management prescription cannot benefit all species. Overall invertebrate species richness and abundance is likely to be promoted by a mosaic of grazing intensities. This can be difficult to achieve on unenclosed upland grassland, though topographical and habitat variation may drive unequal distribution of livestock.

In reality, upland areas are managed for multiple outputs, such as food production, recreation, and ecosystem services, including biodiversity. Within biodiversity management, maintaining specialised or characteristic species or assemblages of upland areas, such as through promoting mosaics of management types and intensities across landscapes, may be seen as a higher priority than maximising species richness, abundance or diversity of any particular taxa. To this end, our findings add to earlier evidence that grazing abandonment may be detrimental to some elements of biodiversity but find no evidence that ground beetle assemblages are detrimentally affected by grazing at a low-intensity compared to at a higher intensity.

## Acknowledgements

We thank the Woodland Trust and their Glen Finglas staff for hosting the grazing experiment and providing logistical support. The work was funded by the Scottish Government’s Rural and Environment Science and Analytical Services’ Strategic Research Programme.

**Supplementary Material 1**: List of all recorded Carabidae species with their full taxonomic name, the subfamily they belong to, and their unique identifier for both the Global Biodiversity Information Facility (GBIF) and for the National Center for Biotechnology Information (NCBI) taxonomy backbones.

**Supplementary Material 2**: Temporal trends in species richness and log abundance of ground beetles at Glen Finglas. Letters A to F represent sampling blocks whilst grazing treatments are shown by numerals I, II, III and IV.

**Supplementary Material 3**: Ecological distances of ground beetle assemblages collected in Glen Finglas. Distances between treatment-years are calculated using Bray-Curtis dissimilarity indices on untransformed mean species abundance matrices. Heatmaps are grouped by sampling blocks. Letters A to F represent sampling blocks whilst grazing treatments are shown by numerals I, II, III and IV.

**Supplementary Material 4**: Results of Simper analysis – species contributing most on the differences between treatments.

**Supplementary material 5**: The effect of treatment and sampling year on the mean abundances of the 16 most common carabid species collected in Glen Finglas. P-values of the linear mixed effect models are shown under each plot, and sampling years are colour coded according to the legend.

## Notes

### Competing Interest Statement

The authors have declared no competing interest.

## Reference

Bassett, E.R.L., Fraser, L.H., 2015. Effects of Cattle on the Abundance and Composition of Carabid Beetles in Temperate Grasslands. Journal of Agricultural Studies 3, 36–47. https://doi.org/10.5296/jas.v3i1.6731

Batáry, P., Báldi, A., Szél, G., Podlussány, A., Rozner, I., Erdős, S., 2007. Responses of grassland specialist and generalist beetles to management and landscape complexity. Diversity and Distributions 13, 196–202. https://doi.org/10.1111/j.l472-4642.2006.00309.x

Blake, S., McCracken, D.I., Eyre, M.D., Garside, A., Foster, G.N., 2003. The relationship between the classification of Scottish ground beetle assemblages (Coleoptera, Carabidae) and the National Vegetation Classification of British plant communities. Ecography 26, 602–616. https://doi.org/10.1034/j.1600-0587.2003.03491.x

Bonari, G., Fajmon, K., Malenovský, I., Zelený, D., Holuša, J., Jongepierová, I., Kočárek, P., Konvička, O., Uřičář, J., Chytrý, M., 2017. Management of semi-natural grasslands benefiting both plant and insect diversity: The importance of heterogeneity and tradition. Agriculture, Ecosystems & Environment 246, 243–252. https://doi.org/10.1016/j.agee.2017.06.010

Brooks, D.R., Bater, J.E., Clark, S.J., Monteith, D.T., Corbett, S.J., Beaumont, D.A., Chapman, J.W., 2012. Large carabid beetle declines in a United Kingdom monitoring network increases evidence for a widespread loss in insect biodiversity. Journal of Applied Ecology 49, 1009–1019. https://doi.org/10.1111/j.1365-2664.2012.02194.x

Brown, G.R., Matthews, I.M., 2016. A review of extensive variation in the design of pitfall traps and a proposal for a standard pitfall trap design for monitoring ground-active arthropod biodiversity. Ecology and Evolution 6, 3953–3964. https://doi.org/10.1002/ece3.2176

Buchanan, G.M., Grant, M.C., Sanderson, R.A., Pearce-Higgins, J.W., 2006. The contribution of invertebrate taxa to moorland bird diets and the potential implications of land-use management. Ibis 148, 615–628. https://doi.org/10.1111/j.1474-919X.2006.00578.x

Butterfield, J., Coulson, J.C., 1983. The carabid communities on peat and upland grasslands in northern England. Holarctic Ecology 6, 163–174.

Clarke, K.R., 1993. Non-parametric multivariate analyses of changes in community structure. Australian Journal of Ecology 18, 117–143.

Coates, P.S., Ricca, M.A., Prochazka, B.G., Brooks, M.L., Doherty, K.E., Kroger, T., Blomberg, E.J., Hagen, C.A., Casazza, M.L., 2016. Wildfire, climate, and invasive grass interactions negatively impact an indicator species by reshaping sagebrush ecosystems. PNAS 113, 12745–12750. https://doi.org/10.1073/pnas.1606898113

Cole, L.J., Morton, R., Harrison, W., McCracken, D.I., D, R., 2008. The influence of riparian buffer strips on carabid beetle (Coleoptera, Carabidae) assemblage structure and diversity in intensively managed grassland fields. Biodiversity Conservation 17, 2233–2245. https://doi.org/10.1007/s10531-007-9304-1

Cole, L.J., Pollock, M.L., Robertson, D., Holland, J.P., Mccracken, D.I., 2006. Carabid (Coleoptera) assemblages in the Scottish uplands⍰: the influence of sheep grazing on ecological structure. October.

Conant, R.T., Cerri, C.E.P., Osborne, B.B., Paustian, K., 2017. Grassland management impacts on soil carbon stocks: a new synthesis. Ecological Applications 27, 662–668. https://doi.org/10.1002/eap.1473

De Cáceres, M., Legendre, P., 2009. Associations between species and groups of sites: indices and statistical inference. Ecology 90, 3566–3574. https://doi.org/10.1890/08-1823.1

De Cáceres, M., Legendre, P., Moretti, M., 2010. Improving indicator species analysis by combining groups of sites. Oikos 119, 1674–1684. https://doi.org/10.1111/j.1600-0706.2010.18334.x

Dennis, P., Skartveit, J., McCracken, D.I., Pakeman, R.J., Beaton, K., Kunaver, A., Evans, D.M., 2007. The effects of livestock grazing on foliar arthropods associated with bird diet in upland grasslands of Scotland. Journal of Applied Ecology 45, 279–287. https://doi.org/10.1111/j.1365-2664.2007.01378.x

Dennis, P., Young, M.R., Howard, C.L., Gordon, I.J., 1997. The response of epigeal beetles (Col.: Carabidae, Staphylinidae) to varied grazing regimes on upland Nardus stricta grasslands. Journal of Applied Ecology 34, 433–443. https://doi.org/10.2307/2404888

Déri, E., Magura, T., Horváth, R., Kisfali, M., Ruff, G., Lengyel, S., Tóthmérész, B., 2011. Measuring the short-term success of grassland restoration: the use of habitat affinity indices in ecological restoration. Restoration Ecology 19, 520–528. https://doi.org/10.1111/j.1526-100X.2009.00631.x

Evans, D.M., Villar, N., Littlewood, N.A., Pakeman, R.J., Evans, S.A., Dennis, P., Skartveit, J., Redpath, S.M., 2015. The cascading impacts of livestock grazing in upland ecosystems: a 10-year experiment. Ecosphere 6, 42. https://doi.org/10.1890/ES14-00316.1

Eyre, M.D., Luff, M.L., Rushton, S.P., Topping, C.J., 1989. Ground beetles and weevils (Carabidae and Curculionoidea) as indicators of grassland management practices. Journal of Applied Entomology 107, 508–517. https://doi.org/10.1111/j.l439-0418.1989.tb00285.x

García, R.R., Jáuregui, B.M., García, U., Osoro, K., Celaya, R., 2009. Effects of livestock breed and grazing pressure on ground-dwelling arthropods in Cantabrian heathlands. Ecological Entomology 34, 466–475. https://doi.org/10.1111/j.1365-2311.2008.01072.x

Gardner, S.M., Hartley, S.E., Davies, A., Palmer, S.C.F., 1997. Carabid communities on heather moorlands in northeast Scotland: The consequences of grazing pressure for community diversity. Biological Conservation 81, 275–286. https://doi.org/10.1016/S0006-3207(96)00148-6

Gardner, S.M., Hartley, S.E., Davies, A., Palmer, S.C.F., 1991. Ground beetle (Coleoptera: Carabidae) communities on upland heath and their association with heathland flora. Journal of Biogeography 18, 281–289. https://doi.org/10.2307/2845398

Gimingham, C.H., 1985. Age-related interactions between Calluna vulgaris and phytophagous insects. Oikos 44, 12–16. https://doi.org/10.2307/3544036

Grandchamp, A.-C., Bergamini, A., Stofer, S., Niemelä, J., Duelli, P., Scheidegger, C., 2005. The influence of grassland management on ground beetles (Carabidae, Coleoptera) in Swiss montane meadows. Agriculture, Ecosystems & Environment 110, 307–317. https://doi.org/10.1016/j.agee.2005.04.018

Hancock, M.H., Legg, C.J., 2011. Pitfall trapping bias and arthropod body mass. Insect Conservation and Diversity no-no. https://doi.org/10.1111/j.1752-4598.2O11.00162.x

Jung, J.-K., Jeong, J.-C., Lee, J.-H., 2019. Effects of pitfall trap size and sampling duration on collection of ground beetles (Coleoptera: Carabidae) in temperate forests. Entomological Research 49, 229–236. https://doi.org/10.1111/1748-5967.12358

Koivula, M., 2011. Useful model organisms, indicators, or both? Ground beetles (Coleoptera, Carabidae) reflecting environmental conditions. ZooKeys 100, 287–317. https://doi.org/10.3897/zookeys.100.1533

Kruess, A., Tscharntke, T., 2002. Contrasting responses of plant and insect diversity to variation in grazing intensity. Biological Conservation 106, 293–302. https://doi.org/10.1016/S0006-3207(01)00255-5

Lengyel, S., Déri, E., Magura, T., 2016. Species richness responses to structural or compositional habitat diversity between and within grassland patches: a multi-taxon approach. PLoS ONE 11, 0149662. https://doi.org/10.1371/journal.pone.0149662

Lindroth, C.H., 1986. The Carabidae (Coleoptera) of Fennoscandia and Denmark, Fauna Entomologica Scandinavica. E.J. Brill, Leiden, Netherlands.

Lindroth, C.H., 1985. The Carabidae (Coleoptera) of Fennoscandia and Denmark. E.J. Brill, Leiden, Netherlands.

Littlewood, N.A., 2008. Grazing impacts on moth diversity and abundance. Society 151–160.

Littlewood, N.A., Pakeman, R.J., Pozsgai, G., 2012. Grazing impacts on Auchenorrhyncha diversity and abundance on a Scottish upland estate. Insect Conservation and Diversity 5, 67–74. https://doi.org/10.1111/j.1752-4598.2011.00135.x

Lövei, G.L., Magura, T., 2011. Can carabidologists spot a pitfall? The non–equivalence of two components of sampling effort in pitfall–trapped ground beetles (Carabidae. Community Ecology 12, 18–22. https://doi.org/10.1556/ComEc.12.2011.1.3

Lövei, G.L., Sunderland, K.D., 1996. Ecology and behavior of ground beetles. Annual Review of Entomology 41, 231–256.

Luff, M.L., 2007. The Carabidae (ground beetles) of Britain and Ireland (Handbooks for the Identification of British Insects). Royal Entomological Society, St. Albans [England].

Luff, M.L., Eyre, M.D., Rushton, S.P., 1992. Classification and prediction of grassland habitats using ground beetles (Coleoptera, Carabidae. Journal of Environmental Management 35, 301–315. https://doi.org/10.1016/S0301-4797(11)80012-5

Luff, M.L., Eyre, M.D., Rushton, S.P., 1989. Classification and Ordination of Habitats of Ground Beetles (Coleoptera, Carabidae) in North-East England. Journal of Biogeography 16, 121. https://doi.org/10.2307/2845086

Luff, M.L., Rushton, S.P., 1989. The ground beetle and spider fauna of managed and unimproved upland pasture. Agriculture, Ecosystems and Environment 25, 195–205. https://doi.org/10.1016/0167-8809(89)90051-0

Lyons, A., Ashton, P.A., Powell, I., Oxbrough, A., 2017. Impacts of contrasting conservation grazing management on plants and carabid beetles in upland calcareous grasslands. Agriculture, Ecosystems and Environment 244, 22–31. https://doi.org/10.1016/j.agee.2017.04.020

Martin, D., Fraser, M.D., Pakeman, R.J., Moffat, A.M., 2013. Natural England Review of Upland Evidence 2012 - Impact of moorland grazing and stocking rates. Natural England Evidence Review Number 006.

Martin, G., Barth, K., Benoit, M., Brock, C., Destruel, M., Dumont, B., Grillot, M., Hübner, S., Magne, M., Moerman, M., Mosnier, C., Parsons, D., Ronchi, B., Schanz, L., Steinmetz, L., Werne, S., Winckler, C., Primih, R., 2020. Potential of multi-species livestock farming to improve the sustainability of livestock farms: A review. Agricultural Systems 181, 102821. https://doi.org/10.1016/j.agsy.2020.102821

McGovern, S., Evans, C.D., Dennis, P., Walmsley, C., McDonald, M.A., 2011. Identifying drivers of species compositional change in a semi-natural upland grassland over a 40-year period. Journal of Vegetation Science 22, 346–356. https://doi.org/10.1111/j.1654-1103.2011.01256.x

Nijssen, M., Alders, K., van der Smissen, N., Esselink, H., 2001. Effects of grass-encroachment and grazing management on carabid assemblages of dry dune grasslands. Proceedings of the Section Experimental and Applied Entomology of the Netherlands Entomological Society (N.E.V.) 12, 113–120.

Oakleaf, J.R., Kennedy, C.M., Baruch-Mordo, S., West, P.C., Gerber, J.S., Jarvis, L., Kiesecker, J., 2015. A world at risk: Aggregating development trends to forecast global habitat conversion. PLoS ONE 10, 0138334. https://doi.org/10.1371/journal.pone.0138334

O’Hara, R.B., Kotze, D.J., 2010. Do not log-transform count data. Methods in Ecology and Evolution 1, 118–122. https://doi.org/10.llll/j.2041-210X.2010.00021.x

Oksanen, J., Blanchet, F.G., Kindt, R., Legendre, P., O’Hara, R.G., Simpson, G.L., Solymos, P., Stevens, M.H.H., Wagner, H., 2010. vegan: Community ecology package.

Pakeman, R.J., Fielding, D.A., Everts, L., Littlewood, N.A., 2019. Long-term impacts of changed grazing regimes on the vegetation of heterogeneous upland grasslands. Journal of Applied Ecology 56, 1794–1805. https://doi.org/10.1111/1365-2664.13420

Pétillon, J., Georges, A., Canard, A., Ysnel, F., 2007. Impact of cutting and sheep grazing on ground – active spiders and carabids in intertidal salt marshes (Western France). Animal Biodiversity and Conservation 2, 201–209.

Pinheiro, J., Bates, D., DebRoy, S., Sarkar, D., R Core Team, 2017. {nlme}: Linear and Nonlinear Mixed Effects Models.

Pozsgai, G., Baird, J., Littlewood, N.A., Pakeman, R.J., Young, M.R., 2018. Phenological changes of the most commonly sampled ground beetle (Coleoptera: Carabidae) species in the UK environmental change network. International Journal of Biometeorology 62, 1063–1074. https://doi.org/10.1007/s00484-018-1509-3

Pozsgai, G., Baird, J., Littlewood, N.A., Pakeman, R.J., Young, M.R., 2015. Long-term changes in ground beetle (Coleoptera: Carabidae) assemblages in Scotland. Ecological Entomology 41, 157–167. https://doi.org/10.1111/een.12288

Pozsgai, G., Littlewood, N.A., 2014. Ground beetle (Coleoptera: Carabidae) population declines and phenological changes: Is there a connection? Ecological Indicators 41, 15–24. https://doi.org/10.1016/j.ecolind.2014.01.029

Pozsgai, G., Littlewood, N.A., 2011. Changes in the phenology of the ground beetle Pterostichus madidus (Fabricius, 1775). Insect Science 18, 462–472. https://doi.org/10.1111/j.1744-7917.2011.01416.x

Pravia, A., Andersen, R., Artz, R.E., Pakeman, R.J., Littlewood, N.A., 2019. Restoration trajectory of carabid functional traits in a formerly afforested blanket bog. Zoologica Academiae Scientiarum Hungaricae 65, 33–56. https://doi.org/10.17109/AZH.65.Suppl.33.2019

Ridding, L.E., Redhead, J.W., Pywell, R.F., 2015. Fate of semi-natural grassland in England between 1960 and 2013: A test of national conservation policy. Global Ecology and Conservation 4, 516–525. https://doi.org/10.1016/j.gecco.2015.10.004

Rodwell, J.S., 1992. Grasslands and montane communities, British Plant Communities. Cambridge University Press, Cambridge, UK.

Rodwell, J.S., 1991. Mires and heaths, British Plant Communities. Cambridge University Press, Cambridge, UK.

Sanderson, R., Newton, S., Selvidge, J., 2020. Effects of vegetation cutting on invertebrate communities of high conservation value Calluna upland peatlands. Insect Conservation and Diversity 13, 239–249. https://doi.org/10.1111/icad.12384

Smith, S.W., Vandenberghe, C., Hastings, A., Johnson, D., Pakeman, R.J., Wal, R., Woodin, S.J., 2014. Optimizing carbon storage within a spatially heterogeneous upland grassland through sheep grazing management. Ecosystems 17, 418–429. https://doi.org/10.1007/s10021-013-9731-7

Söderström, B., Svensson, B., Vessby, K., Glimskär, A., 2001. Plants, insects and birds in semi-natural pastures in relation to local habitat and landscape factors. Biodiversity and Conservation 10, 1839–1863. https://doi.org/10.1023/A:1013153427422

Suominen, O., Niemelä, J., Martikainen, P., Niemelä, P., Kojola, I., 2003. Impact of reindeer grazing on ground-dwelling Carabidae and Curculionidae assemblages in Lapland. Ecography 26, 503–513. https://doi.org/10.1034/j.1600-0587.2003.03445.x

Thomas, C.F.G., Brown, N.J., Kendall, D.A., 2006. Carabid movement and vegetation density: Implications for interpreting pitfall trap data from split-field trials. Agriculture, Ecosystems and Environment 113, 51–61. https://doi.org/10.1016/j.agee.2005.08.033

Török, P., Valkó, O., Deák, B., Kelemen, A., Tóth, E., Tóthmérész, B., 2016. Managing for species composition or diversity? Pastoral and free grazing systems in alkali steppes. Agriculture, Ecosystems and Environment 234, 23–30. https://doi.org/10.1016/j.agee.2016.01.010

Török, P., Valkó, O., Deák, B., Kelemen, A., Tóthmérész, B., 2014. Traditional cattle grazing in a mosaic alkali landscape: effects on grassland biodiversity along a moisture gradient. PloS one 9, e97095. https://doi.org/10.1371/journal.pone.0097095

Tóth, E., Deák, B., Valkó, O., Kelemen, A., Miglécz, T., Tóthmérész, B., Török, P., 2018. Livestock type is more crucial than grazing intensity: traditional cattle and sheep grazing in short-grass steppes. Land Degradation and Development 29, 231–239. https://doi.org/10.1002/ldr.2514

Twardowski, J.P., Pastuszko, K., Hurej, M., Gruss, I., 2017. Effect of different management practices on ground beetle (Coleoptera: Carabidae) assemblages of uphill grasslands. Polish Journal of Ecology 65, 400–409. https://doi.org/10.3161/15052249PJE2017.65.3.007

WallisDeVries, M.F., Poschlod, P., Willems, J.H., 2002. Challenges for the conservation of calcareous grasslands in northwestern Europe: Integrating the requirements of flora and fauna. Biological Conservation 104, 265–273. https://doi.org/10.1016/S0006-3207(01)00191-4

Wang, L., Delgado-Baquerizo, M., Wang, D., Isbell, F., Liu, Jun, Feng, C., Liu, Jushan, Zhong, Z., Zhu, H., Yuan, X., Chang, Q., Liu, C., 2019. Diversifying livestock promotes multidiversity and multifunctionality in managed grasslands. Proc Natl Acad Sci USA 116, 6187–6192. https://doi.org/10.1073/pnas.1807354116

York, A., 2000. Long-term effects of frequent low-intensity burning on ant communities in coastal blackbutt forests of southeastern Australia. Austral Ecology 25, 83–98. https://doi.org/10.1046/j.1442-9993.2000.01014.x

