## Supplementary figures and images for "Grazing impacts on ground beetle (Coleoptera: Carabidae) abundance and diversity on a semi-natural grassland"

### Supplementary Material 2

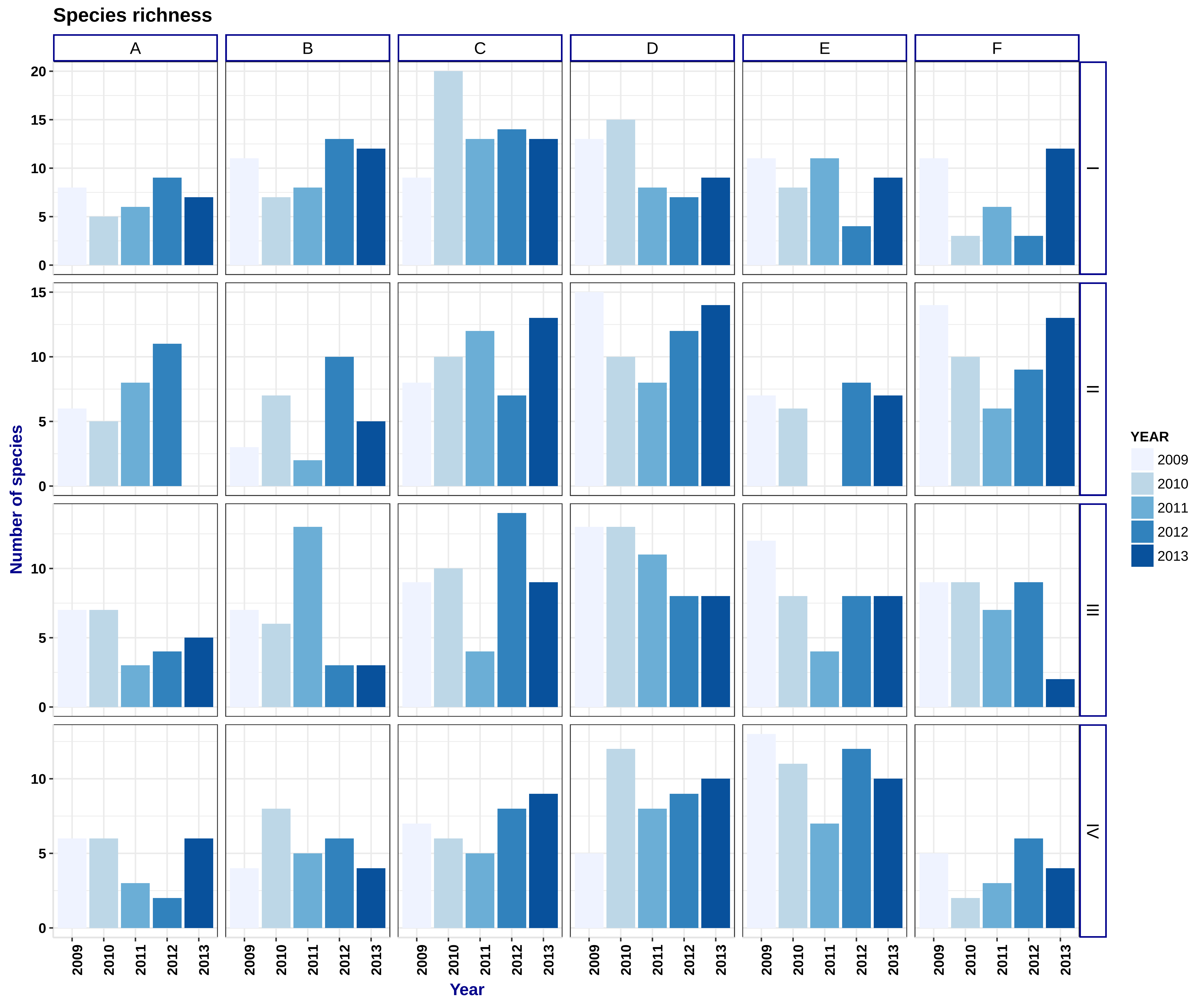

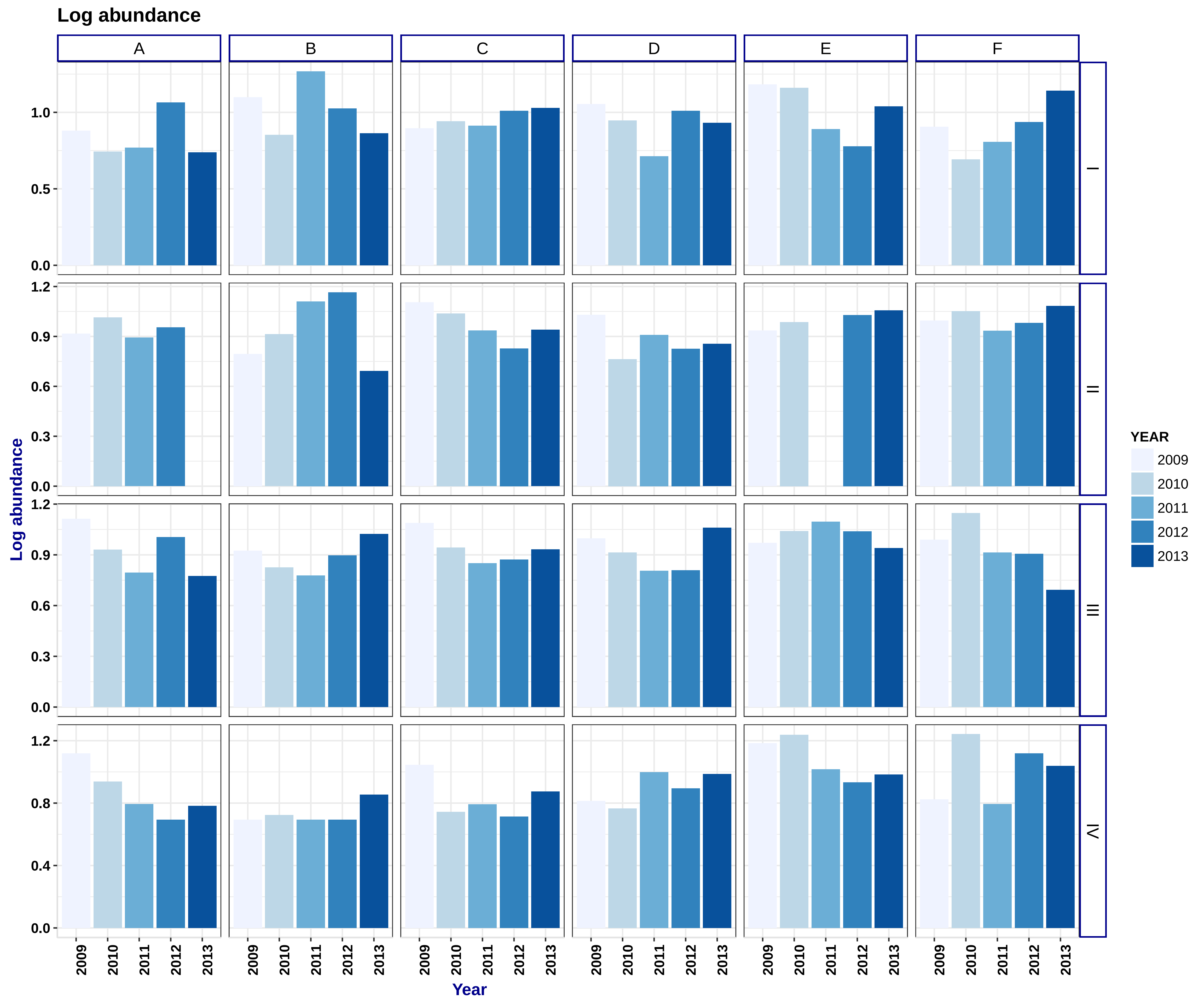

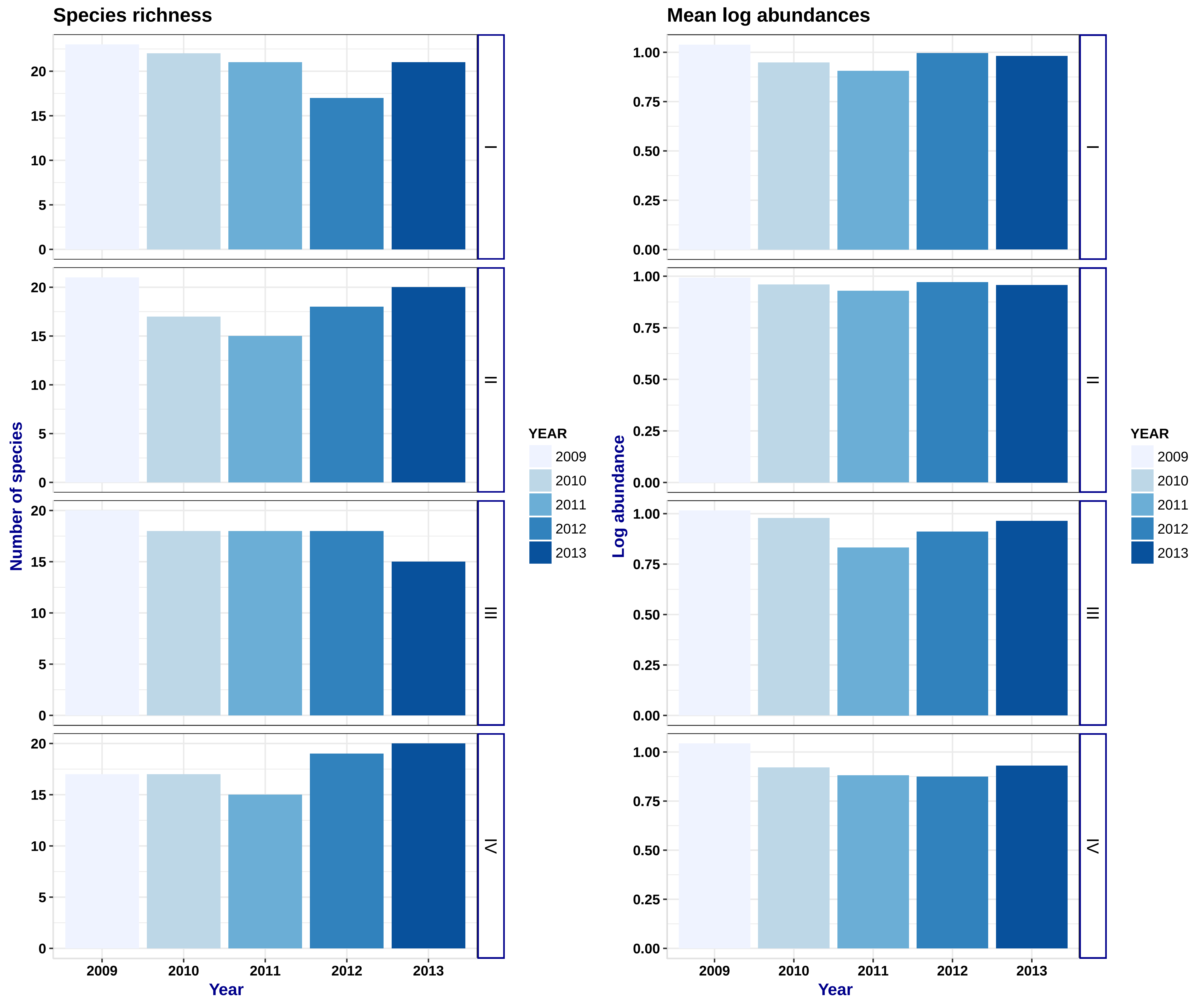

### Supplementary Material 3

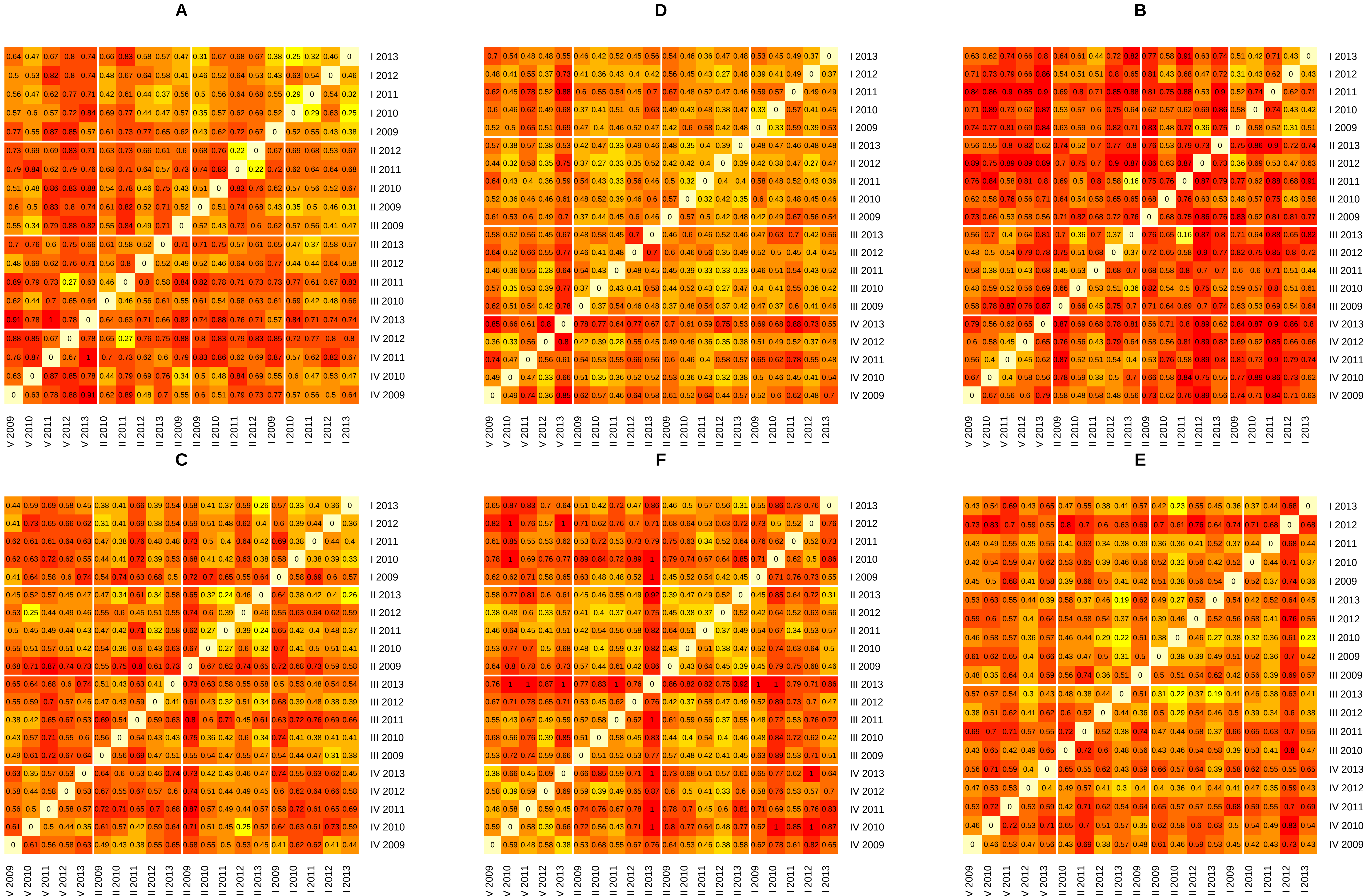

### Supplementary Material 5

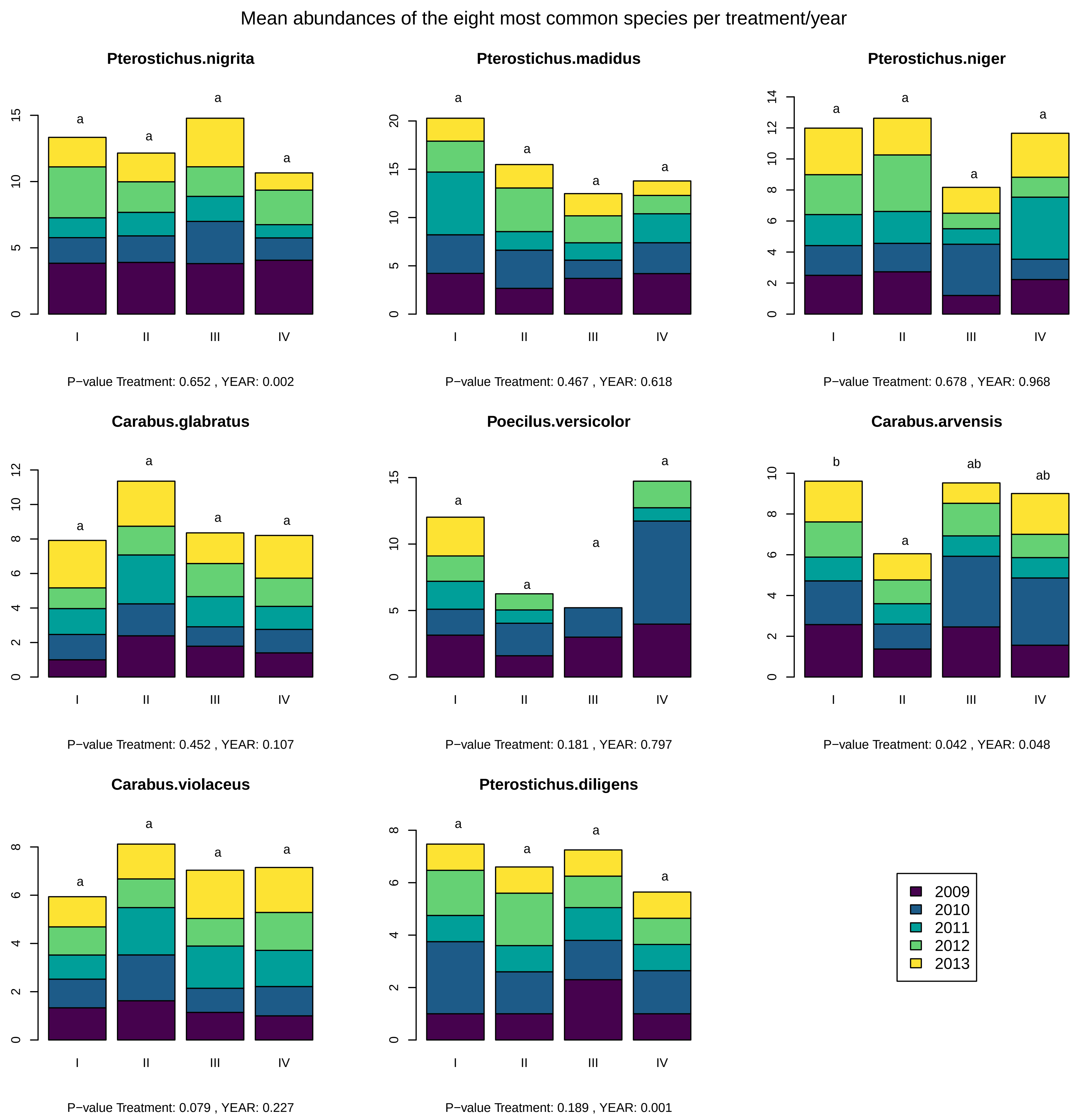

Mean abundances of the eight most common species per year/treatment

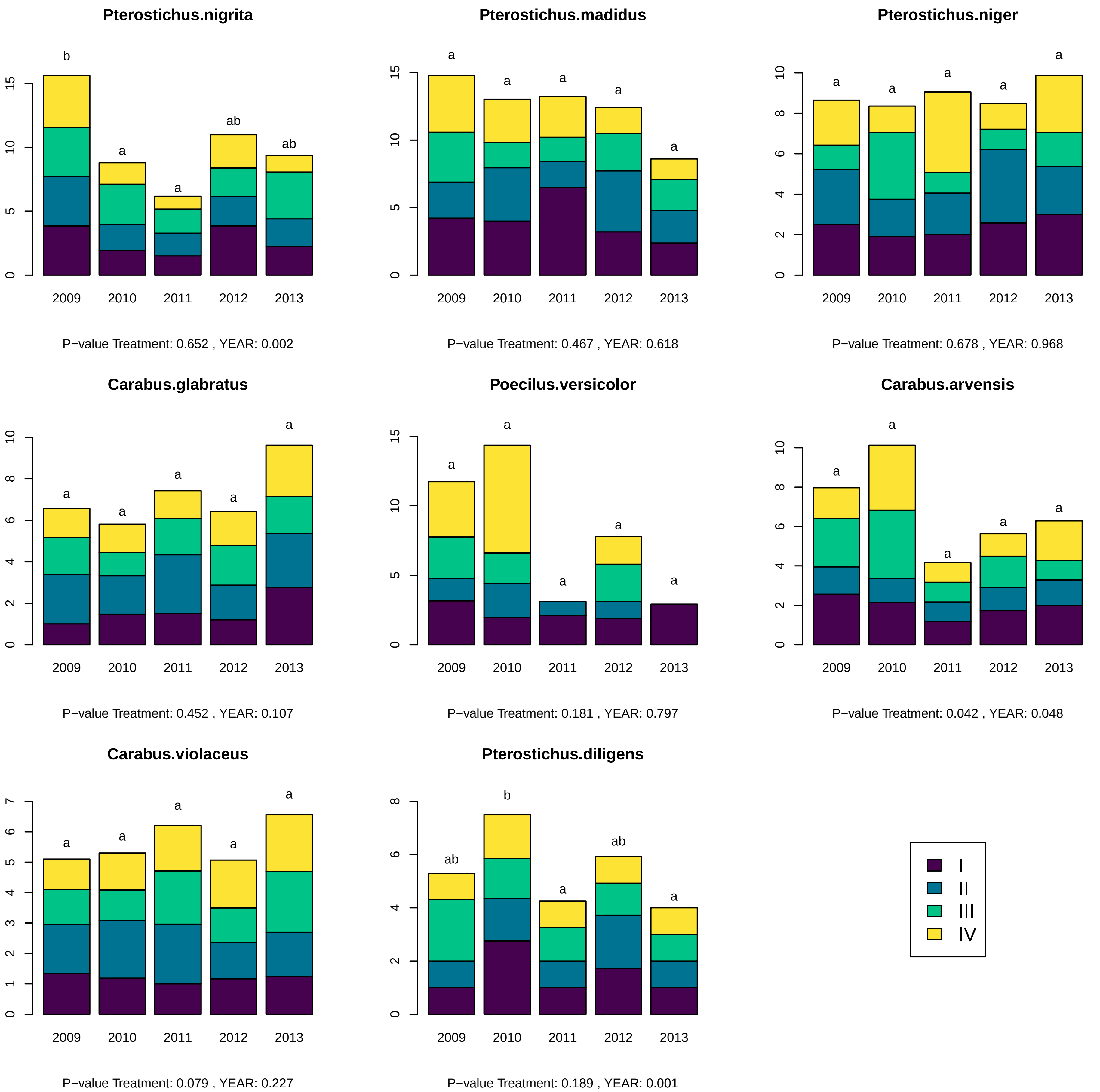
